# Single-cell DNA methylation sequencing by combinatorial indexing and enzymatic DNA methylation conversion

**DOI:** 10.1101/2022.10.09.511389

**Authors:** Zac Chatterton, Praves Lamichhane, Diba Ahmadi Rastegar, Lauren Fitzpatrick, Hélène Lebhar, Christopher Marquis, Glenda Halliday, John B Kwok

## Abstract

DNA methylation is a critical molecular mark involved in cellular differentiation and cell-specific processes. Single-cell whole genome DNA methylation profiling methods hold great potential to resolve the DNA methylation profiles of individual cell-types. Here we present a method that couples single-cell combinatorial indexing (sci) with enzymatic conversion (sciEM) of unmethylated cytosines. The method facilitates single-base resolution DNA methylation profiling of single-cells that is highly correlated with single-cell bisulfite-based workflows (r^2^ >0.99) whilst improving sequencing alignment rates, reducing adapter contamination and over-estimation of DNA methylation levels (CpG and non-CpG). As proof-of-concept we perform sciEM analysis of the temporal lobe, motor cortex, hippocampus and cerebellum of the human brain to resolve single-cell DNA methylation of all major cell-types.

## Introduction

The covalent addition of a methyl group to cytosine bases in mammalian DNA (DNA methylation) is one of the most highly studied epigenetic modifications (1). Primarily occurring in the CpG context, DNA methylation is critical for organism development (2) and plays an essential role in regulating gene expression during cellular differentiation (3). Cell-types have highly specific DNA methylation patterns (4) necessitating the analysis of DNA methylation in pure cellular populations, however limited cell surface markers or highly interconnected tissue networks prohibit cell isolation from tissues such as the human brain.

Single-cell whole genome bisulfite sequencing techniques have recently been described (5-8) that can produce single-base resolution DNA methylation information from which cell-specific whole genome DNA methylation profiles (methylomes) can be reconstructed bioinformatically. However, these sequencing library preparations are prohibitively expensive for most labs because of high reagent costs associated with single-cell single-well reactions. Recently a single-cell combinatorial indexing (sci-) bisulfite sequencing approach (termed sciMET) was described in which nuclei are sorted and tagged with sequencing indexes over multiple rounds, forming unique combinations of indexes per nuclei (9). The sciMET approach allows multiple-nuclei single-well reactions, thus reducing reagent costs.

Whole Genome Bisulfite Sequencing (WGBS) is the gold standard method for DNA methylation analysis (10, 11) but is not without limitations. It has been estimated that 84–96% of DNA is degraded during the bisulfite conversion reaction (12). Additionally, methylated DNA is overrepresented in WGBS libraries leading to an over-estimate of DNA methylation levels (13) particularly in CHG and CHH contexts (14). This is especially relevant in the analysis of DNA methylation in embryonic stem cells and neurons that have been reported to exhibit high levels of CHG and CHH methylation (15-20).

Unmethylated cytosines can also be deaminated by *APOBEC* enzymes, resulting in base changes analogous to bisulfite conversion (sequenced as T) (21). Notably, enzymatic conversion is less degradative to DNA and can produce high quality single-base resolution DNA methylation data (21). Such attributes may be particularly beneficial in situations where DNA content is limited such as the single-cell analysis of DNA methylation. However, enzymatic conversion in single cells is challenging due, in part, to multiple reaction cleanup steps required that results in DNA loss. A major advantage of the sci-approach over single-cell/single-well methods is the ability to perform deamination reactions of multiple cells per-well, thus increasing per-well DNA content. Here we combine sci-with enzymatic conversion (termed sciEM, Figure 1a) and show application by characterizing single-base DNA methylation profiles of human brain cell-types without the need for cell-type markers (eg. NeuN). The sciEM approach accurately captures CpG methylation dynamics across annotated regulatory features of the human genome. Both CpG and non-CpG (CpH) methylation estimates are lower than bisulfite conversion and we find no evidence of higher global CpH DNA methylation within neurons from the temporal lobe, motor cortex, hippocampus or cerebellum of the human brain. The sciEM approach represents an economical method for single-cell single-base resolution DNA methylation analysis.

**Figure 1.**
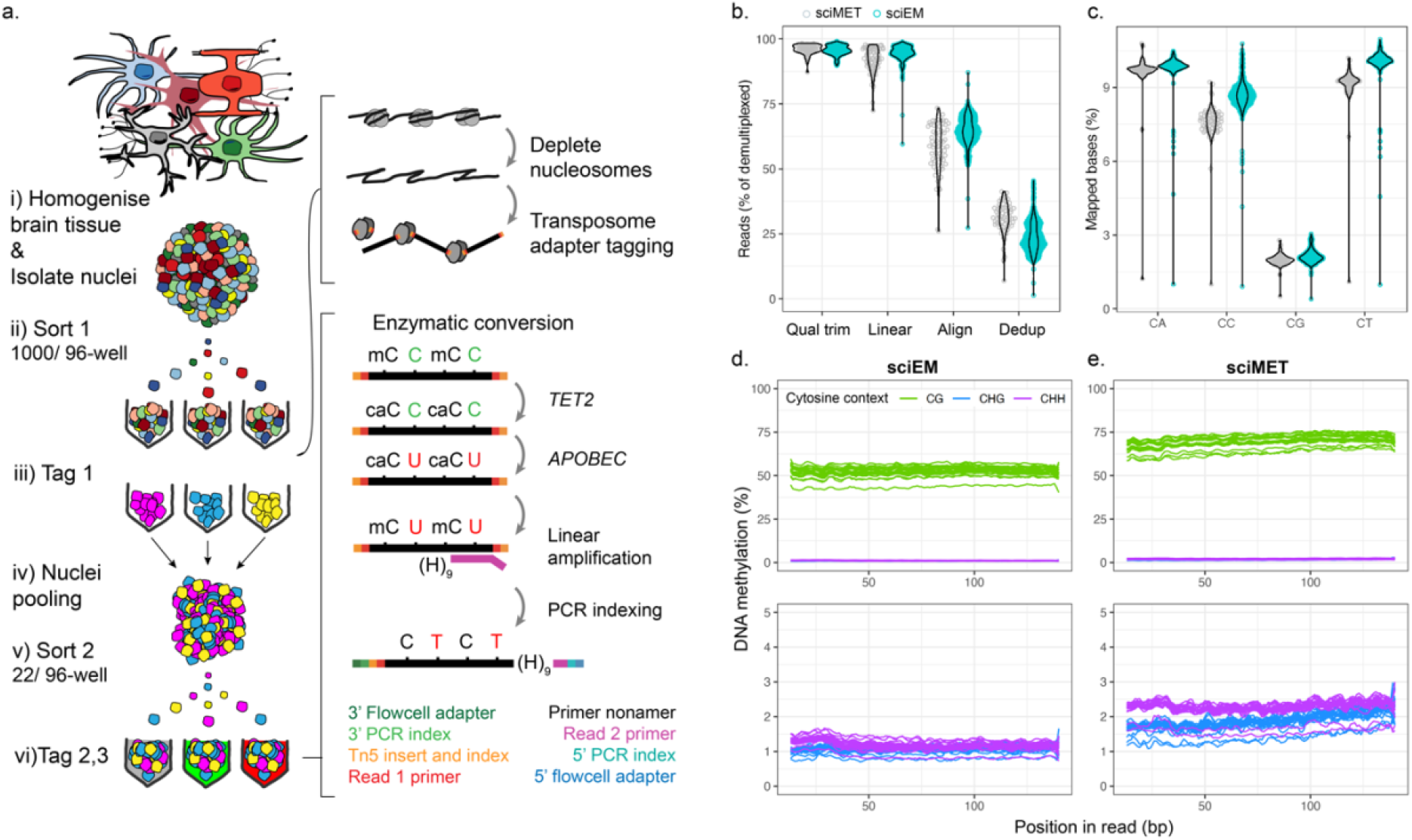
a. Single-cell combinatorial indexing and enzymatic conversion (sciEM) workflow in which i) whole tissue (eg. brain tissue) is homogenized to dissociates cells. Nuclei from heterogeneous cell-types are isolated and ii) sorted by Fluorescent Activated Nuclei-Sorting (FANS). iii) Nuclei membranes are permeabilized, nucleosomes depleted, and molecular tags (tag 1, Tn5 barcode) are attached to genomic DNA via transposome tagmentation. iv) Nuclei are pooled, vi) re-sorted by FANS and vi) unmethylated cytosines are converted to thymine following treatment with *TET2* and *APOBEC* enzymes and Linear amplification. Molecular tags 2,3 (i5 & i7 barcodes) and sequencing adapters are attached via PCR amplification (adapted from (9)). b. Per cell read processing metric’s. c. Cytosine dinucleotides covered as percentage of mapped bases. d & e DNA methylation bias plots for sciEM and sciMET methods respectively for reads mapping to autosome’s and the X-chromosome, with close-up of CHG and CHH DNA methylation (bottom). H=A, C or T.

## Results

### Library construction and sequencing read processing

Using frozen post-mortem brain tissue from mouse (NextSeq, n=1) and human (NextSeq, n=1 & NovaSeq, n=4) we were able to construct both sciMET and sciEM single-cell libraries in parallel. Within the sciEM workflow we use a G-depleted (mg) random linear primer that we observed to improve CpH mapping within preliminary experiments (NextSeq, Supplementary Figure 1). Both bisulfite and enzymatic conversion efficiencies were high, 99.99% and 99.94% respectively, however the sciMET method produced ∼10X the amount of library than sciEM (518nM v 53nM by RT-qPCR). Post-sequencing (NovaSeq), single-nuclei were identified by unique barcode combinations (Tn5, i5 & i7 barcodes). Single-nuclei with >100 unique mapped reads were observed to have significantly higher mapping efficiency, more paired-reads, larger insert sizes and a lower proportion of reads mapped using local alignment (Students t-test p-value’s < 5×10^−17^, Supplementary Figure 1a-d.), representing high-quality single-nuclei. Following k-means clustering of unique mapped reads, a total of 710 and 64 high quality single-nuclei from sciEM and sciMET workflows passed were retained for analysis (Supplementary Figure 1e-f), representing 54 and 58% of the nuclei fluorescently sorted for each workflow respectively. The mapping efficiency was 58 ± 5% and 64.9 ± 10% for sciMET and sciEM libraries respectively (Figure 1b.), but we note the sciMET mapping efficiencies were lower than previously reported (9), results that are partially attributed to the removal of 4% of reads that contained substantial linear primer sequences (Figure 1b.). The number of mapped reads were higher in sciMET (mean 254,194 ±191,560) than sciEM (150,515 ± 147,388) (Students t-test p-value = 7.25×10^−5^, Figure 1b) leading to higher coverage of mappable cytosine dinucleotides (e.g., CpG p=1.91×10^−10^, Supplementary Figure 3a). However, the library loading concentration largely influences read counts and, proportional to mapped reads, the sciEM method covered a greater number of all cytosine dinucleotides, particularly CpT and CpC dinucleotides (p < 6.1×10^−10^, Figure 1c).

### DNA methylation

Global DNA methylation levels were observed to be lower in sciEM than sciMET libraries for CpG (54% v 71%), CHG (1.0 v 1.2%) and CHH (1.2 v 2.2%) cytosine contexts (p-value < 7.10×10^−8^). WGBS library preparation methods have previously been reported to have an over-representation of methylated fragments (13). In line with these reports, we observed an increase in the 5mC levels from 5’ to 3’ of sciMET reads for CpG, CHG and CHH contexts (student t-test p-value < 0.003, first 50% bases v last 50% bases). In contrast, 5mC levels were stable across sciEM reads (Figure 1d.). Of note, the 5mC levels of chromosome 21 were significantly lower than other chromosomes, an effect that was pronounced within sciEM libraries. We observed a very high correlation between the CpG methylation of sciEM and sciMET across annotated genomic features (e.g., R^2^=0.996 +/-5kb Ensemble genes, Pearson’s p-value=6.9×10^−^ _120_).

CpG hypomethylation is generally associated with chromatin accessibility and gene activity and sciEM successfully captured global DNA methylation dynamics across regulatory regions e.g. hypomethylation of gene promoter regions, CpG Islands and open chromatin (DNase-seq) (Figure 2a-c.). We observed CpG hypomethylation across annotated regions enriched for active histone marks (e.g. H3K4me3, H3K9ac, H3K27ac) and H3K4me1 boundaries (Figure 2d-g). The H3K27me3 histone modification is a marker of bivalent polycomb regulated promoters in which dynamic crosstalk between DNA methylation controls gene expression (22) and we observed CpG hypomethylation across genomic regions enriched for H3K27me3 (Figure 2h). Conversely, hypermethylation was observed within annotated regions enriched for repressive histone modifications (H3K36me3 and H3K9me3) (Figure 3i-j). Of note, sciEM CpG methylation levels are significantly lower than sciMET measurements e.g., 14% lower across annotated genes (+/-5kb) (paired t-test p-value= 3.46×10^−65^). The difference in CpG DNA methylation between sciEM and sciMET was highly correlated to the underlying CpG methylation levels (Figure 2k).

**Figure 2.**
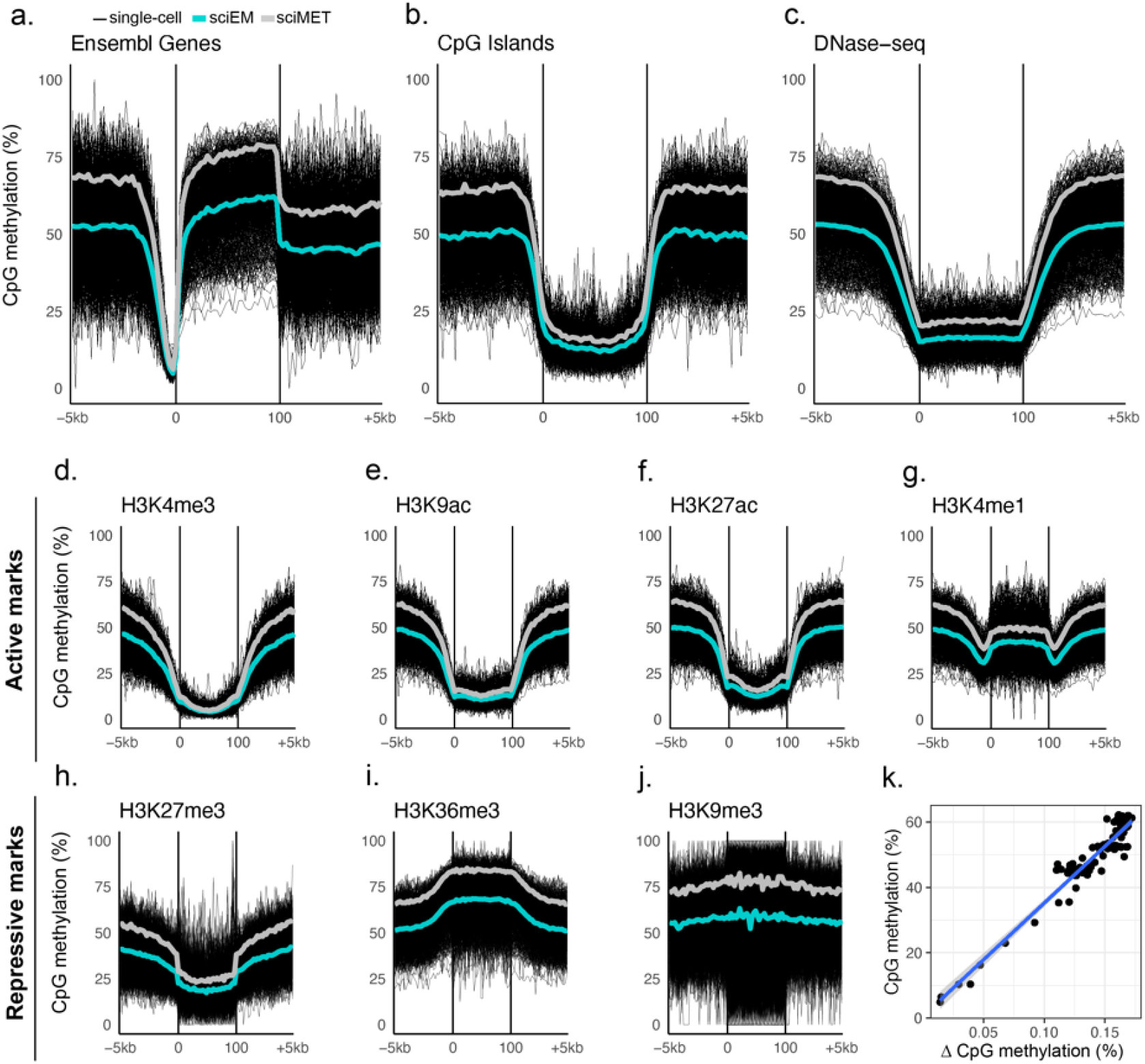
CpG methylation across genomic features. The mean CpG methylation levels of single-cells (black) from sciEM (cyan) and sciMET (grey) protocols across a. Ensemble genes b. CpG Islands and regions of c. DNase hypersensitivity as well as histone modifications associated with active (d-g) and repressive (h-j) chromatin conformations. k) Scatterplot of CpG methylation levels (sciMET mean) and methylation difference (sciMET-sciEM) across each bin (3%) of annotated Ensemble genes (+/-5kb).

**Figure 3.**
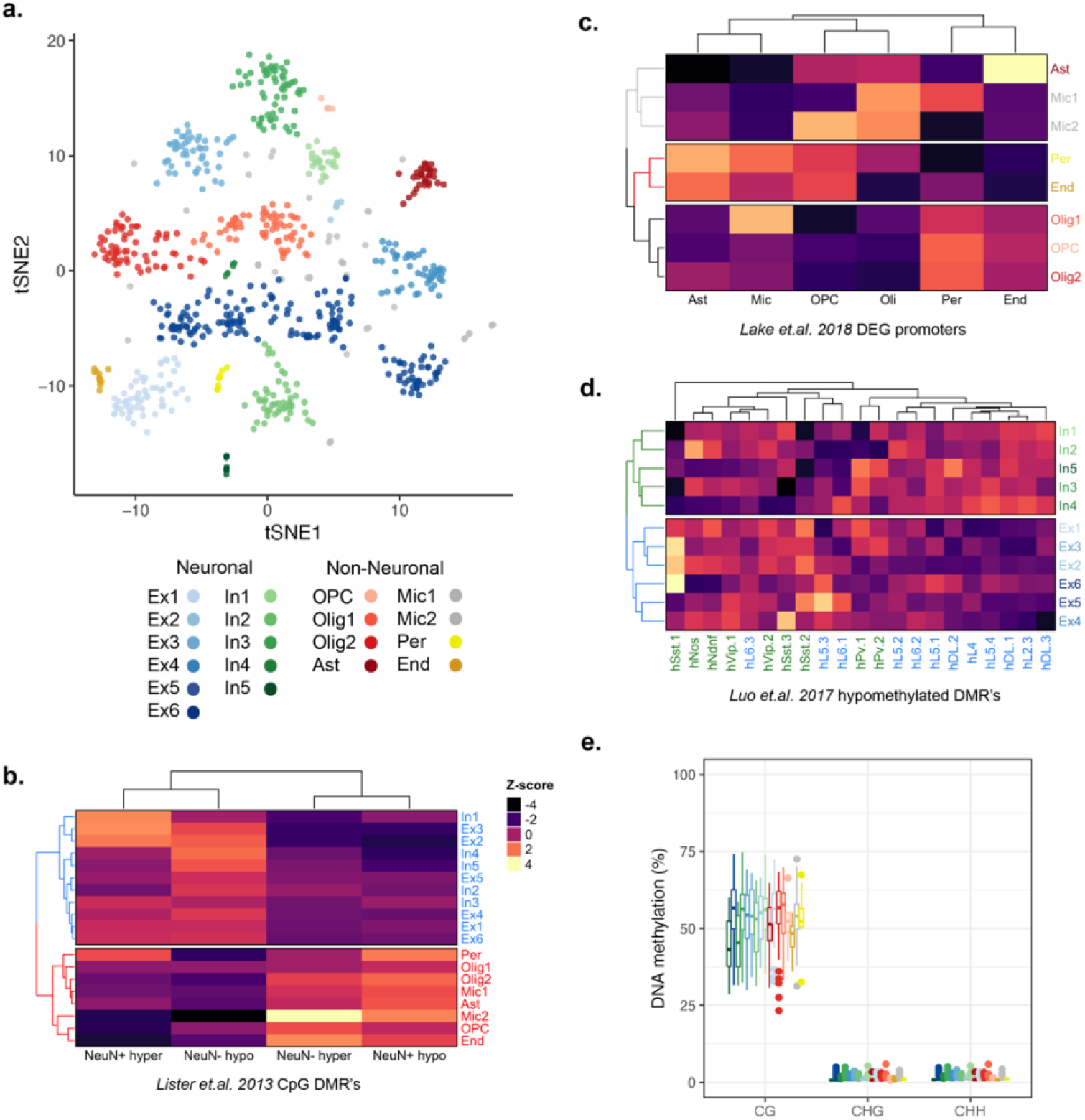
Cell-type discrimination by sciEM single nuclei DNA methylation. a. Single nuclei DNA methylation clustering (NMF-tSNE). Clusters (n=19) are defined by unique colors. b. Heatmap of summarized CpG methylation z-score’s of clusters across annotated Neuron and Non-neuronal DMR’s. c. Heatmap of summarized CpG methylation z-score’s of non-neuronal cell clusters across annotated non-neuronal DEG’s. d. Heatmap of summarized CpG methylation z-score’s of neuronal clusters across annotated neuronal subtype CpG DMR’s. e. Boxplots of CpG, CHG and CHH DNA methylation of each cell-type.

### Single-cell clustering

To assess the ability of sciEM to discriminate cell types we summarized CpG DNA methylation of regulatory regions (Ensembl Regulatory Build) and CpH methylation across 100kb genomic bins (used to cluster Neuronal cell-types (7)) across 710 high quality single nuclei (methods). The DNA methylation information of each single nuclei was combined using NMF and projected into 2-dimensional space (tSNE) from which 19 clusters were identified (Figure 3a). We observed distinct patterns of DNA methylation within established Differentially Methylated Regions (DMRs) distinct to Neurons and Non-Neuronal cell-types from the human brain (17) which enabled the identification of 11 neuron (n=430) and 8 non-neuronal clusters (n=280) (Figure 3b). As gene promoter hypomethylation is associated with gene activity, the promoter DNA methylation status of established Differentially Expressed Genes (DEG’s) of non-neuron cell-types (23) enabled the identification of Astrocytes (n=30), Endothelial cells (n=12), Microglia (n=45), Oligodendrocyte precursor cells (OPC’s, n=5), Oligodendrocytes (n=130) and Pericytes (n=8) (Figure 3c). Using the CpG DNA methylation levels at annotated neuronal cell-type CpG DMRs (7) we identified 6 excitatory (n=279) and 5 inhibitory (n=151) neuronal cell-type clusters (Figure 3d). We found no significant difference in the per-nuclei CpG, CHG or CHH DNA methylation levels between neuron and non-neuronal cell-type clusters (Figure 3e).

## Discussion

To our knowledge we present the first enzyme based single-cell DNA methylation method with single-base resolution. The sciEM method extends single-cell combinatorial indexing approaches developed using sodium bisulfite (sciMET). Bisulfite sequencing is problematic for single-cell sequencing as it degrades the limited amount of DNA in each cell (24), however enzymatic based conversion of unmethylated cytosines has been shown to be less degradative to the DNA, resulting in more genomic coverage, even at 100 pg amounts (21). To generate sciEM libraries we used a G-depleted random primer (linear amplification primer) that binds more efficiently to genomic fragments devoid of cytosines following conversion (enzymatic or bisulfite converts ∼95% of cytosines to uracils). G-depleted random primers have been previously shown to improve library complexity, coverage uniformity and reduce artefactual reads (8). Generally, the sciEM approach results in higher library loss during construction, and a lower amount of library input into the second barcoding PCR (i5 & i7), preferential amplification of smaller molecules (reduced insert size), and a higher rate of duplicate sequences compared to sciMET. However, the data quality is high. We show high correlation of DNA methylation between sciEM and sciMET (r^2^=0.99) approaches, accurate recapitulation of DNA methylation dynamics across gene features and the ability to resolve single base resolution of single-cell types.

Since the first whole genome bisulfite sequencing (WGBS) study, a multitude of techniques have been developed to characterize genome-wide methylation (25). However, as DNA methylation patterns are unique to single-cell-types, it is essential to move towards single-cell DNA methylation profiling (26). In disease states, DNA methylation patterns are known to be altered, leading to aberrant cascades of molecular changes (3). Hence it is important to have accurate base level estimates of DNA methylation levels. Whole genome bisulfite sequencing approaches have been reported to over-represent methylated fragments (13). We observed 5mC bias within sciMET sequencing reads, elevating from 5’ to 3’, that were not observed within sciEM. Further, 5mC levels were lower in sciEM, an effect that was greater within CHH loci (typically unmethylated) relative to CpG loci (typically methylated). Whilst we cannot rule out over conversion of 5mC within sciEM, our observations of a 5mC bias within sciMET sequencing reads (elevating from 5’ to 3’), comparatively higher 5mCHH (typically unmethylated) then 5mCpG (typically methylated) and a slightly higher conversion efficiency indicate a potentially over-representation of methylated fragments in sciMET.

Neurons exhibit distinct DNA methylation patterns, particularly in non-CpG (CpH) loci, compared to non-neuronal brain cells (17, 27) as well as between neuronal subtypes (7) (28). To our knowledge we present the first whole genome DNA methylation assessment of brain cell-types using enzymatic DNA methylation assessment. We did not observe significantly higher CpH methylation within neurons as previously reported (17). Further, our single-cell analysis of brain cell types omits the use the NeuN antibody for neuron selection, hence we cannot rule out the possibility that NeuN-positivity (nuclei surface marker) of Neurons relates to CpH DNA methylation.

The brain is a highly heterogeneous environment, comprising multiple neuronal and glial cell types with unique functions and at various stages of differentiation. Previously, the understanding of disease mechanisms progressed by studying cell populations in bulk which revealed only the average features of the population’s constituents and can obscure the cell-to-cell variability. Since the first whole genome bisulfite sequencing (WGBS) study, a multitude of techniques have been developed to characterise genome-wide methylation (25) at single-base resolution. Moreover, as DNA methylation patterns are unique to cell-types, it is essential to move towards single-cell DNA methylation profiling (26). Single-cell strategies have yielded novel mechanistic insights into brain function (29). The sciEM method represent’s an economical, high-throughput alternative to bisulfite to analyse single-cell DNA methylation at single-base resolution.

## Methods

### Transposome production

Recombinant transposase enzyme (Tn5) was grown (pTXB1-Tn5 vector) and purified following the protocols described in Picelli et al. (30). Cytosine depleted sciMET transposase-loaded oligonucleotides (1-96) were annealed (10 μL each 100 μM) to 10 μL 5′-[Phos]-CTGTCTCTTATACACATCT-3’ oligonucleotide (100 μM) within 80 μL EB buffer (Qiagen), incubating 2 min at 95°C and cooled to room temperature (0.1°C/sec), following protocols (31). Annealed oligonucleotides were dilute 2:5 (EB buffer), mixed with glycerol (50% final solution) and loaded (equal volume) into the recombinant Tn5 (15uM) by incubation for 20 min at room temperature. Annealed oligonucleotide loading was confirmed by gel-shift assay and fragmentation efficiency of each transposome was confirmed (>50%) by qPCR analysis (32).

### Brain sample and nuclei isolation

NextSeq – Post-mortem flash-frozen prefrontal cortex tissue was obtained from a 93-year-old female donor with no diagnosis of neurological disease. Post-mortem flash-frozen cortex was obtained from a genetically modified (*C9orf72*) mouse. Following the protocols of Mulqueen et al, Brain tissue sections were resuspended in 5 mL of ice-cold NIB-HEPES solution (20mM HEPES, 10mM NaCl, 3mM MgCl_2_). The tissues were equilibrated (5 min) and then dounce homogenized (10 loose strokes and 5 tight strokes) and filtered through 35-40μm cell strainers (BD Biosciences, 352235). Nuclei were pelleted (600g) and were transferred to a fresh tube containing 5 mL ice cold NIB-HEPES solution.

NovaSeq - Post-mortem flash-frozen tissue from the Primary Motor Cortex (BA4), Banks of the superior temporal sulcus (BA 22,41/42 [BA22]), Cerebellum (CRB) and Hippocampus (HIP) were obtained from a 47-year-old female donor with no diagnosis of neurological disease. Brain tissue was acquired from the NeuroBioBank (NIH) and approved by the Research Integrity and Ethics Administration of the University of Sydney. A high amount of cellular debris was observed by nuclei isolation protocols described in Mulqueen et al (9), therefore nuclei were isolated instead following protocols described in Matevossian et al (33) and resuspended in 5 mL ice cold NIB-HEPES solution.

### Nucleosome depletion

Following the protocols of Mulqueen et al, nuclei were cross-linked using 135 μL of 37% formaldehyde, quenched with 400 μL of 2.5M glycine and resuspended in 5 mL of ice-cold NIB (10mM Tris HCl pH 7.4, 10 mM NaCl, 3 mM MgCl_2_, 0.1% Igepal (v/v), 1x protease inhibitors) solution, pelleted (500g for 5 min), and washed using 900 μL of 1x NEBuffer 2.1 (NEB, B7202). To denature proteins, nuclei were mixed with 800 μL of 1x NEBuffer 2.1 and 12μL SDS solution (20%) and incubated at 42°C for 30 min with vigorous shaking. Nuclei were then mixed with 20 μL of 10% Triton-X (Sigma, 9002-93-1) and incubated at 42°C for 30 min solubilize proteins/ increase nuclei permeabilization.

### Fluorescent Activated Nuclei-Sorting (FANS) and Tagmentation

The nuclei were stained using 8 μL of 5mg/mL DAPI dye (Thermo-Fisher, Cat. D1306) and filtered through a 35-40μm cell strainer. FANS was performed on BD InFlux-7L (sort 1), separating 1000 single nuclei per well in a 96-well plate containing 5 μL of 2xTB buffer (20 mM Tris(hydroxymethyl)aminomethane, 10 mM MgCl2 and 20% (v/v) dimethylformamide (DMF)) and 5 μL of NIB solution. To each well, 4 μL of 4.56 μM unique transposome (1-96) was added and incubated at 55°C for 15 min with gentle shaking, adding the “Tn5 index”. All wells were then pooled, re-stained with 8 μL of 5 mg/mL DAPI and filtered. FANS was performed again (sort 2), separating 22 or 10 (control wells) single nuclei per well in a 96-well plate containing 2.5 μL of M-digestion buffer (Zymo, Cat. D5020-9), 0.25 μL of Proteinase K (Zymo, D3001-2-5), and 2.5μL of H_2_O. Nuclei were then digested at 50°C for 20 min with gentle shaking and the plate was then spun at 600g for 5 min at 4°C.

### Bisulfite conversion

Prior to bisulfite conversion, 35 pg of pre-tagmented unmethylated lambda DNA was spiked into wells receiving 10 nuclei (sort 2). NextSeq; Each well was made up to 50uL with H_2_O and bisulfite conversion was performed following manufacturer protocols using the EZ-96 DNA Methylation Kit (Zymo, Cat. D5004) and eluted twice (12.5 uL each using elution buffer) for a final volume of 25uL. NovaSeq; Each well was made up to 20 uL with H_2_O and bisulfite conversion was performed following manufacturer protocols using the MethylCode BC conversion kit (Applied Biosystems, Cat. MECOV50) and eluted twice (12.5 uL each using elution buffer) for a final volume of 25uL

### Enzymatic DNA methylation (EM) conversion

Prior to the EM conversion, 35 pg of pre-tagmented unmethylated lambda DNA and 14 pg of the tagmented CpG methylated pUC19 DNA were spiked into wells receiving 10 nuclei (sort 2). Sample volumes were made up to 20uL with H_2_O and cleanup was performed using 1.8X using AMPure XP cleanup beads (Beckman Coulter, Cat. A63881) following manufacturer protocol with the exception that samples were incubated for 10mins at room temperature followed by a single wash step using 60uL 80% EtOH and eluted 29uL of elution buffer. Enzymatic conversion was then performed using the NEBNext Enzymatic Methyl-seq Conversion Module (New England Biolabs, Cat. E7125L) following manufacturers protocol (steps 1.5 to 1.9.11) for inserts 370–420 bp. Briefly, 5-Methylcytosines and 5-Hydroxymethylcytosines were oxidized using TET2 enzyme. DNA was cleaned using AMPure XP cleanup beads in place of NEBNext Sample Purification Beads. DNA was denatured in 0.1 N NaOH and cytosines were deaminated by APOBEC3A, cleaned and eluted using 25 μL of Elution Buffer.

### Linear amplification

Full elution’s from both the bisulfite converted and EM converted libraries were transferred to a plate prepared with the following: 16 μL of PCR-clean H2O, 5μL of 10xNEBuffer 2.1, 2μL of 10mM dNTP mix (New England Biolabs, Cat. N0447), and 2 μL of 10μM of either the 9-nucleotide random primer (n9) previously described in the sciMET protocols (9) or the G-depleted (mg) random primer (8), containing a partial Illumina Standard Read 2 sequencing primer 5′-GGAGTTCAGACGTGTGCTCTTCCGATCT(H1:33340033)(H1)(H1)(H1)(H1)(H1)(H1)(H 1)(H1)-3’. Four rounds of linear amplification were performed using 10U of Klenow (3’-5’ exo) polymerase (Enzymatics, Cat. P7010-LC-L) followed by AMPure XP cleanup (1.1X) and elution in 21μL of 10mM Tris-HCl (pH 8.5) as previously described (9).

### Library indexing and quantification

Indexing PCRs were performed in a 96-well plate to incorporate i5 and i7 indexes. The full elution’s from the linear amplification reaction were mixed with 2 μL each of the 10μM forward and reverse indexing primers (9), 25 μL of 2xKAPA NEBNext Q5 Hot Start HiFi PCR Master Mix (New England Biolabs, Cat. M0543L), and 0.5μL of 100x SYBR Green I dye (FMC BioProducts, Cat. 50513). Real-time PCR was performed on a QuantStudio 6 Flex real-time thermocycler (Applied Biosystems) with the following thermocycling conditions: 95 °C for 2 min, 20 cycles of 94°C for 80 s, 65 °C for 30 s and 72 °C for 30 s [image]. The libraries were then pooled, cleaned using AMPure XP beads (0.8X) and eluted in 20 μL of 10mM Tris-HCl (pH 8.5) as previously described (9). Quantification of each sciMET(n9), sciMET(mg) and sciEM(combined n9 & mg) were performed using the KAPA qPCR Illumina library quantification kit (Kapa Biosystems Cat. KR0405) and the mean of each sciMET result (n9 = 397nM, and mg=638nM) was calculated.

### Library sequencing

NextSeq - sciMET(n9), sciMET(mg) and sciEM (combined n9 and mg libraries) were quantified separately by High Sensitivity D1000 ScreenTape (Agilent, Cat. 5067-5584). Libraries were pooled and sequenced on the Illumina NextSeq 500 (v2 2 × 75 bp cycle Mid-Output Kit) using a 0.9 pM loading concentration, 30% PhiX and custom Read 1 and Index 2 (i5) oligonucleotides matching chemistry temperatures (9). Sequencing was performed using custom chemistry (Read1: 100 imaged cycles; Read2: 10 imaged cycles; Index1: 10 imaged cycles; Index2: 11 imaged cycles, 9 dark cycles, and 9 imaged cycles).

NovaSeq - sciMET(n9) and sciEM(mg) libraries were pooled and then quantified (as above). Libraries were sequenced on the NovaSeq 6000 (v1.5 2 × 300 bp SP Kit) using 116 pM loading concentration, 10% PhiX and custom Read 1 (as above). Sequencing was performed using custom chemistry (Read1: 142 imaged cycles; Index1: 10 imaged cycles; Index2: 7 dark cycles, 10 imaged cycles, 16 dark cycles, and 11 imaged cycles; Read2: 142 imaged cycles).

### Bioinformatics

All scripts used for the processing and analysis of sciMET/sciEM data have been deposited and documented within https://github.com/zchatt/scimet_scripts.

### Sequence read demultiplexing

NextSeq – BCL files were converted to fastq format using bcl2fastq “--create-fastq-for-index- reads --with-failed-reads --use-bases-mask Y*,I10,I20,Y*” generating 2 Read files (R1[100bp] & R2[25bp]) and 2 Index files (I1[10bp, i7 index] & I2[20bp; Tn5 & i5 indexes]) for each sequencing lane. Each R1, R2, I1 and I2 from multiple sequencing lanes were combined by linux cat command and I2 was split into individual Tn5[11bp] and i5[9bp] index files using linux awk command. Fastq files were demultiplexed if all 3 indexes (i5, i7 and Tn5) had a Hamming distance < 3 from the reference, as previously described (9).

NovaSeq - BCL files were converted to fastq format using bcl2fastq “--create-fastq-for-index- reads --use-bases-mask Y*,I10,I21,Y*” generating 2 Read files (R1[142bp] & R2[142bp]) and 2 Index files (I1[10bp, i7] & I2[21bp; Tn5 & i5]) for each sequencing lane. Reads were processed as described above with the exception I2 was split into individual Tn5[11bp] and i5[10bp] index files.

### Sequence read trimming, alignment and DNA methylation extraction

Reads were firstly trimmed (trim 1) using TrimGalore! Software (v0.38.0) with options “-- illumina --stringency 3”. A high number of sequences corresponding to the Linear Primer and read-through of the P7 flow-cell were observed, therefore reads were trimmed again (trim 2) using cutadapt software (v1.8.3) with options “-- anywhere=AGATCGGAAGAGCACACGTCTGAACTCCAGTCA -- anywhere=GAAGAGCACACGTCTGAACTC -- anywhere=ATCTCGTATGCCGTCTTCTGCTTGAAAAAAAAAAGGGGGGGGGGGGGGGGGGGGGGGGGGGG --minimum-length=20 --times=2” (34). Read 2 sequences from the NovaSeq 6000 instrument displayed increasing “G” content >60bp that were largely poly-G sequences (1.2% reads) indicative of low signal intensity. Read 2 sequences were truncated using fastp software (v0.19.6) with options “--max_len1 60”. The human (GRCh38) or mouse (GRCm39) reference genomes were each combined with the lambda phage reference genome that is used for bisulfite/enzymatic conversion control. Alignment of reads were performed using scBS-map software using the options “-l 9 -p 12 -n 10” (35). Aligned reads were deduplicated using samtools software with options “rmdup” (36). DNA methylation information was extracted from aligned deduplicated BAM files using cgmaptools with options “convert bam2cgmap” (37).

### Single-cell discrimination and Quality Control

Single nuclei with < 100 unique mapped reads were removed. The unique read counts of single nuclei have previously been used to discriminate high quality single cells (9). Briefly, k-means clustering (k=3) of unique aligned reads per barcode (k-means, k=3) was performed and normal distributions were fitted to each cluster (Supplementary Figure 2.). Barcodes with unique read counts passing 95% confidence interval threshold (cluster 1) were retained (64 sciMET & 710 sciEM). Bisulfite and enzymatic conversion efficiencies were, calculated as the 5mC % of reads aligned to unmethylated lambda phage genome. Mapping efficiencies were assessed (reads aligned / reads assigned per barcode). We determined the number of uniquely mappable cytosine dinucleotides by intersecting the within the hg38 reference genome with umap files (k=100) downloaded https://bismap.hoffmanlab.org/ (38) using bedtools software with options “getfasta” and umap files (k=100). NextSeq reads were processed as above with the exception that no unique mapped read thresholds were applied, and single nuclei were assigned as mouse or human based on the greatest read mapping efficiency to gr39 and hg38 genomes independently.

### DNA methylation across genomic annotations

CpG and CpH methylation were summarized (3% window) for each single-cell across (+/-5 kb) Ensembl gene annotations, CpG Islands (CGI) as well as ChIP-seq and DNase-seq annotations from the middle frontal cortex (ENCFF146VKE, ENCFF225RTW, ENCFF600AYY, ENCFF724XKK, ENCFF727KZF, ENCFF729EZH, ENCFF835ZYG, ENCFF860MVH from https://www.encodeproject.org/) using cgmaptools with options “mfg” (37). Frontal gyrus NeuN+/- CpG Differentially Methylated Regions (DMRs) generated by Lister et al (17) were downloaded from http://brainome.ucsd.edu/BrainMethylomeData/CG_DMR_lists.tar.gz and converted to hg38 using rtracklayer and hg18ToHg38.over.chain. Neuron CpG DMRs for each of the 21 Neuron cluster described by Luo et al. (7) were converted to hg38 using rtracklayer and hg19ToHg38.over.chain. The hg38 locations of Differentially Expressed Genes (DEG’s) across non-neuron cell-types identified by Lake et al. (23) were extracted using R software biomart package (v 2.46.3) and were separated into gene body and promoter (1.5kb upstream TSS). CpG and CpH methylation were summarized for each DMR and DEG across each single-cell using cgmaptools with options “mtr” (37).

### Cell-type clustering analysis

We performed non-negative matrix factorization (NMF) on summarized CpH methylation across 100kb genomic bins and CpG methylation across the Ensembl Regulatory Build (39) setting k=12, as previously described (9). CpH and CpG NMF matrices were weighted, merged by cell, and plotted into two-dimensional space using students t-distributed stochastic neighbor embedding (t-SNE). Cell clustering was performed using DBSCAN, as previously described (9) using an epsilon value of 1.3. Clustering analysis was performed using all sciEM single-cells (n=710) identifying 16 clusters (Figure 3A). In addition, we evaluated clustering using both sciMET and sciEM single-cells using summarized CpH and CpG (Supplementary Figure 4), summarized CpG alone (Supplementary Figure 5), as well as summarized CpG for sciEM single-cells alone (Supplementary Figure 6). To identify the cell-type of each cluster, sequencing reads of all cell-types within a cluster were collapsed and CpG methylation was summarized for NeuN+/- DMR’s, as described above. Broad subtypes of non-neuronal cells were further classified by CpG methylation summarization of non-neuron DEG promoters, as described above. Non-neuron cell subtypes were defined by lowest (hypomethylated promoters) z-score (annotation × cluster matrix). Broad excitatory and inhibitory neuron subtypes were classified by CpG methylation summarization of promoter CpG DMRs of 21 neuron subtypes, as described above, and defined by hierarchical clustering of z-scores. We performed a linear regression analysis between neuron (n=430) and non-neuronal (n=280) cell-types using per-nuclei CpG, CHG and CHH DNA methylation levels controlling for read depth using R statistic software.

## Declarations

### Ethics approval and consent to participate

The use of human brain tissue was approved by the Research Integrity and Ethics Administration of the University of Sydney (project number: 2018/861). The use of a genetically modified (*C9orf72*) mouse was approved by the Garvan Animal Ethics Committee (project number: 16_14).

### Consent for publication

Not applicable

## Acknowledgements

We thank R.Lister for sharing plasmid vectors; R.Mulqueen, R.Lister, J.Pflueger and S. Freytag for discussions throughout the project; S.Pineda for sharing mouse tissue for the pilot studies.

## Availability of data and materials

The dataset(s) supporting the conclusions of this article are available in the SRA repository, [currently processing, will be available prior to publication].

## Authors’ contributions

Z.C conceived the study and coordinated experiments. G.H performed post-mortem brain tissue dissections. Z.C, P.L, D.A.R and L.F performed nuclei isolation, fluorescent activated nuclei sorting and sequencing library construction. Z.C performed bioinformatic and statistical analysis. H.L and C.M performed recombinant protein and isolation experiments. Z.C and J.B.K wrote the manuscript, with contributions from all authors. All authors read and approved the final manuscript.

