## Supplementary Material for "Single-cell DNA methylation sequencing by combinatorial indexing and enzymatic DNA methylation conversion"

**NextSeq500 sequencing of sciEM and sciMET libraries using regular (n9) and modified (mg) linear primers**

We discriminated single-nuclei using k-means clustering of unique mappable reads (described in methods), in which 7, 12, 158 and 295 single-nuclei generated from sciEM(mg), sciEM(n9), sciMET(mg) and sciMET(n9) library preparation methods passed threshold (95% CI). The small number of single cells passing threshold limited our ability to contrast mouse and human nuclei, however provided sufficient information to evaluate read quality between library preparation methods. We observed significantly fewer reads removed in sciEM libraries (0.6% mean) compared to sciMET (12.5%) due to quality issues (Students p-value = $1.67 \times 10^{-164}$, Supplementary Figure 1a). We observed a high proportion of sciMET library reads that were contaminated with linear primer sequences (methods), particularly within single cells prepared using the sciMET(mg) library preparation method (74% of quality trimmed reads). Following quality trimming and linear primer removal, the alignment efficiencies were similar between sciEM(mg), sciEM(n9) and sciMET(n9) libraries. We observed a significantly higher coverage of mappable CpH sites within the sciEM(mg) than sciEM(n9) (Students t-test p-value = 0.03). No difference was detected between sciEM(mg) and sciMET(n9) libraries in coverage of mappable CpG or CpH (Students t-test p-value > 0.14). We also observed typical high CpG and low CpH DNA methylation within the sciMET(mg) as well as the sciMET(n9) libraries, giving us confidence in the DNA methylation output.

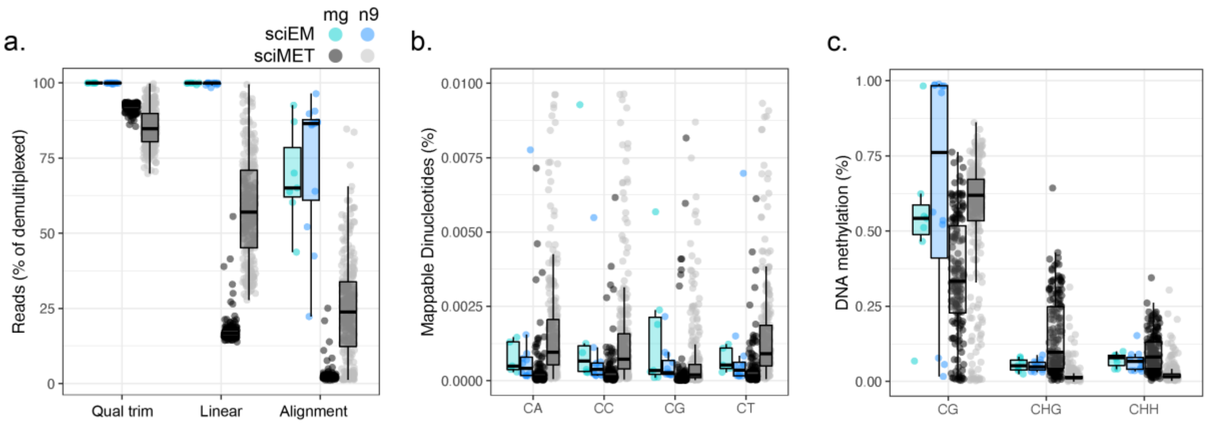


Supplementary Figure 1. Read quality evaluations of sciEM and sciMET libraries sequenced on NextSeq500 platform. Boxplots of a. read processing metrics b. cytosine dinucleotides covered as percentage of mapped bases, and c. CpG and CpH DNA methylation levels.

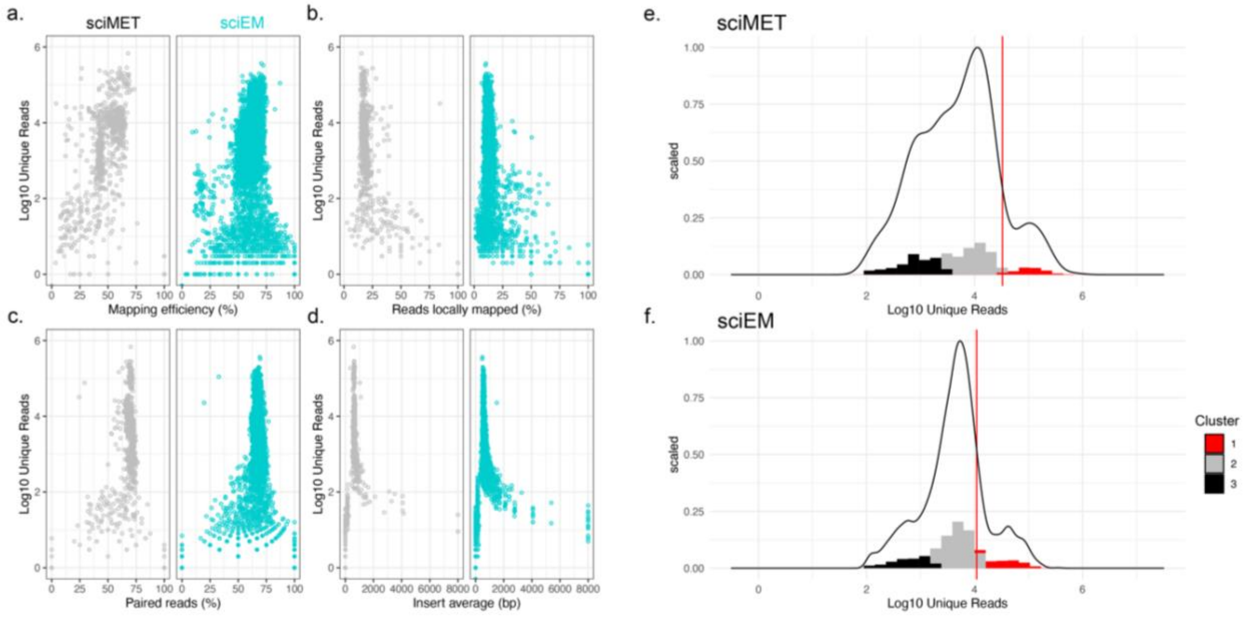


Supplementary Figure 2. Single-cell characterization using unique read thresholding. Scatter plots showing the log10 unique reads for each detected barcode (Tn5, i5 & i7 combined) compared to a) mapping efficiency, b) reads mapped by local alignment strategy, c) paired reads, and d) average insert size. Density plots and histograms of unique reads from barcodes (>100 unique reads) expected from e) sciMET libraries and f) sciEM libraries. Histograms are colored by k-means cluster and 95% CI cluster 1 (red line).

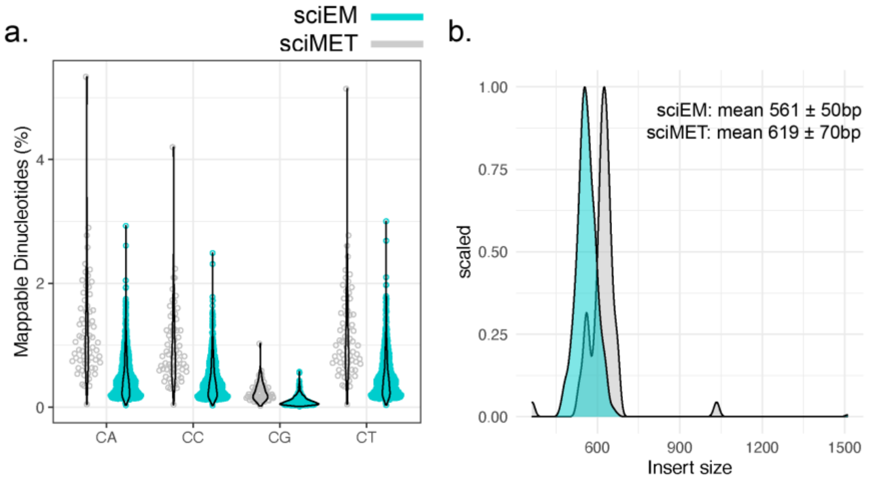


Supplementary Figure 3. Mapping and read features of sciEM and sciMET libraries. a. Genome-wide mapped of cytosine dinucleotides. b. Histogram of insert size lengths.

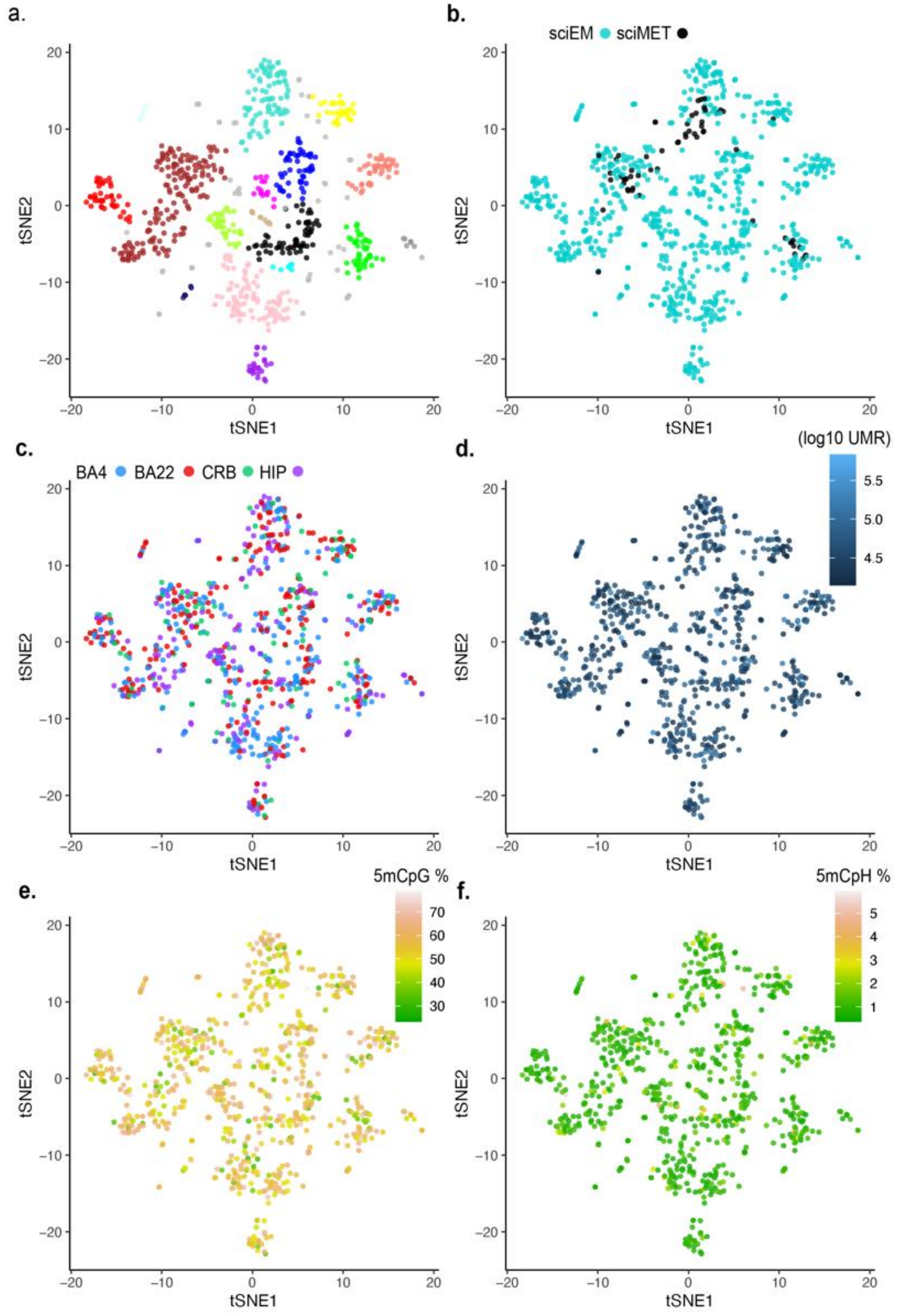


Supplementary Figure 4. Clustering of sciEM (n=710) and scMET (n=64) single nuclei using summarized CpH (100kb bins) and CpG regulatory DNA methylation. a. Cell clusters (n=19) were identified using dbscan algorithm following NMF-tSNE (methods) and re-colored by b. the sequencing method. c. the brain region in which the nuclei were derived. d. the number of Unique Mapped Reads (UMR) sequenced for each single nuclei. e. global CpG DNA methylation percentage, and f. global CpH DNA methylation percentage.

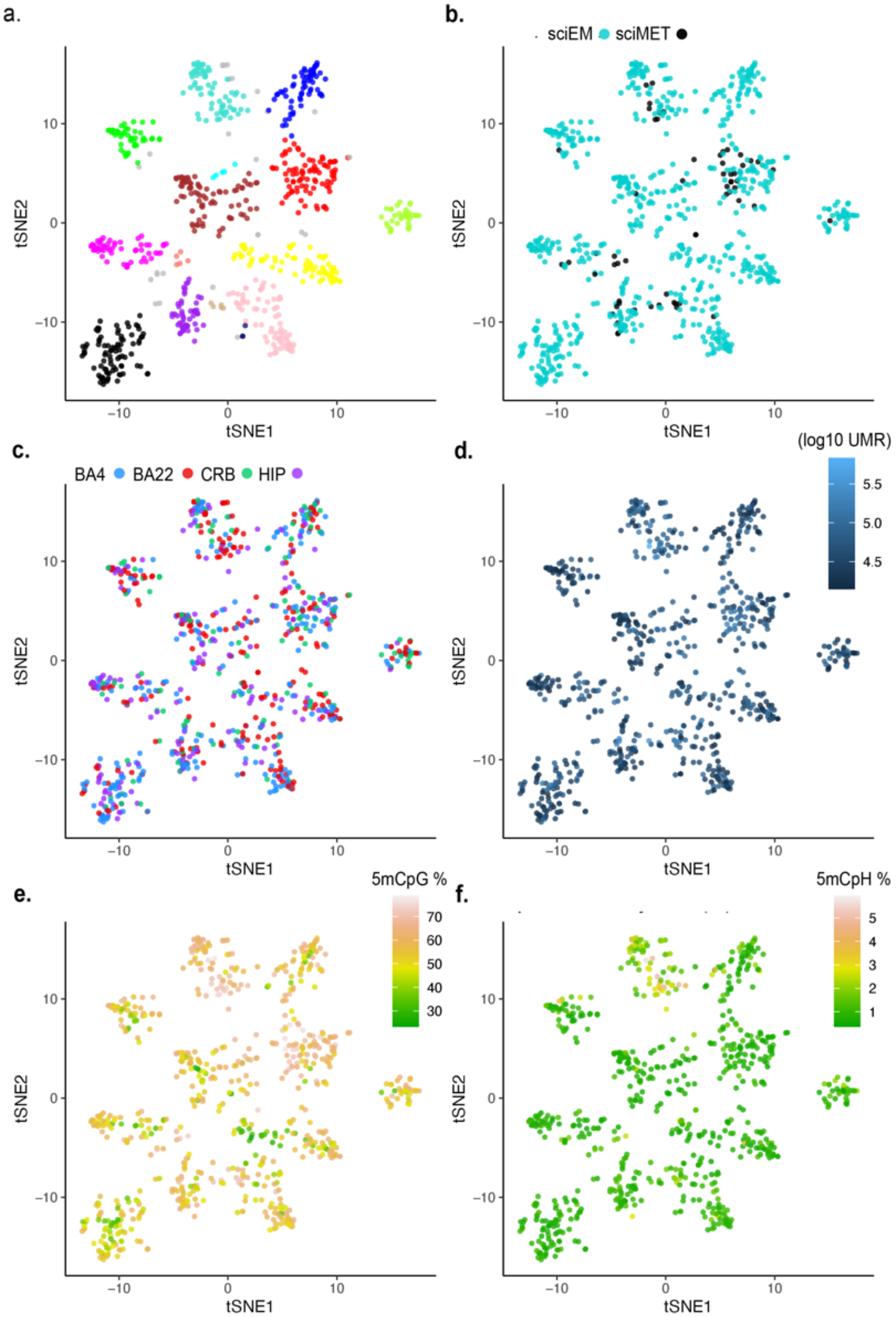


Supplementary Figure 5. Clustering of sciEM (n=710) and scMET (n=64) single nuclei using summarized CpG regulatory DNA methylation. a. Cell clusters (n=16) were identified using dbscan algorithm following NMF-tSNE (methods) and re-colored by b. the sequencing method. c.

the brain region in which the nuclei were derived. d. the number of Unique Mapped Reads (UMR) sequenced for each single nuclei. e. global CpG DNA methylation percentage, and f. global CpH DNA methylation percentage.

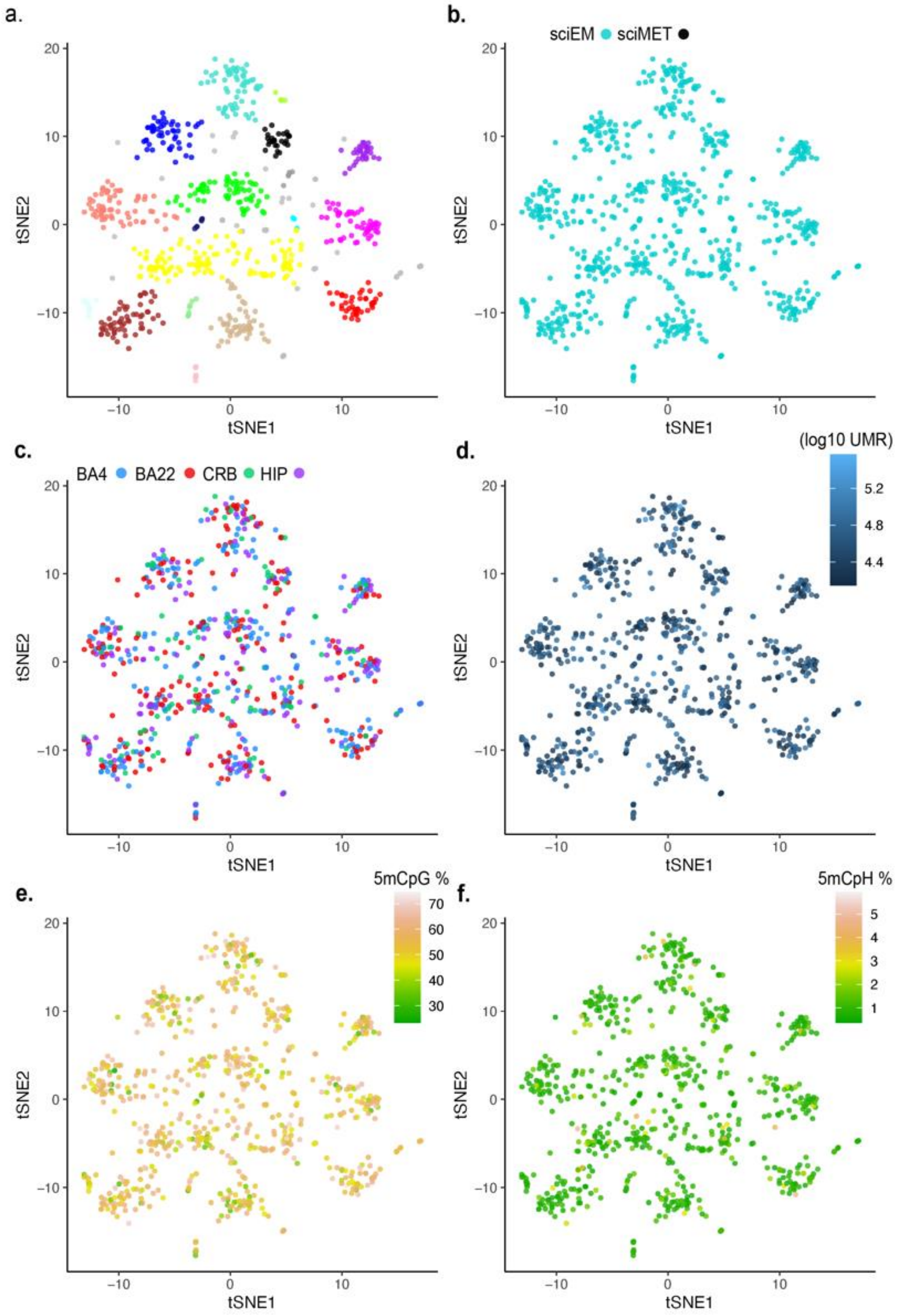


Supplementary Figure 6. Clustering of sciEM (n=710) single nuclei using summarized CpG regulatory DNA methylation. a. Cell clusters (n=18) were identified using dbscan algorithm following NMF-tSNE (methods) and re-colored by b. the sequencing method. c. the brain region in which the nuclei were derived. d. the number of Unique Mapped Reads (UMR) sequenced for each single nuclei. e. global CpG DNA methylation percentage, and f. global CpH DNA methylation percentage.
